# Highly efficient genome modification of cultured primordial germ cells with lentiviral vectors to generate transgenic songbirds

**DOI:** 10.1101/2020.08.11.245753

**Authors:** Ivana Gessara, Falk Dittrich, Moritz Hertel, Staffan Hildebrand, Alexander Pfeifer, Carolina Frankl-Vilches, Mike McGrew, Manfred Gahr

## Abstract

The ability to genetically manipulate organisms has led to significant insights in functional genomics in many species. In birds, manipulation of the genome is hindered by the inaccessibility of the one-cell embryo. During embryonic development, avian primordial germ cells (PGCs) migrate through the blood stream and reach the gonadal *anlage*; where they develop into mature germ cells. Here, we explored the use of PGCs to produce transgenic offspring in the zebra finch, which is a major animal model for sexual brain differentiation, vocal learning and vocal communication. Zebra finch PGCs (zfPGCs) obtained from embryonic blood significantly proliferated when cultured in an optimized culture medium and conserved the expression of germ and stem cell markers. Transduction of cultured zfPGCs with lentiviral vectors was highly efficient leading to strong expression of the enhanced green fluorescent protein (eGFP). Transduced zfPGCs were injected into the host embryo and transgenic songbirds were successfully generated.

## INTRODUCTION

Songbirds are major animal models to study the genetic and neural basis of vocal learning and communication (Mooney, 2020; Mello, 2014; Prather et al., 2017) as well as sex hormone dependent brain development (Gahr, 2007; Balthazart et al., 2010; McCarthy and Arnold, 2011) and adult neurogenesis (Goldman and Nottebohm, 1983; Paton and Nottebohm, 1984). For the zebra finch, transgenic models have been successfully developed (Agate et al., 2009; Abe et al., 2015; Liu et al., 2015). Agate et al. (2009) were the first to inject lentiviral vectors for GFP into the blastodisc of freshly laid zebra finch eggs in order to target the primordial germ cells (PGCs), the precursors of spermatocytes and oocytes. However, due to inefficiency of the method, only two other transgenic models have been generated over the past 10 years using this method (Abe et al., 2015; Liu et al., 2015).

During avian development, PGCs are located in the central area of the blastodisc until Eyal-Giladi and Kochav (EGK) stage X (Eyal-Giladi et al., 1976; Kochav et al., 1980); from here, PGCs translocate to the germinal crescent (Ginsburg et al., 1986) and finally migrate through the developing vascular system to reach the gonadal *anlage* (Fujimoto et al., 1976; Nakamura et al., 1988; Jung et al., 2019). Avian PGCs are easily isolated from the circulatory system by blood aspiration between Hamburger and Hamilton (HH) stages 14 and 17 (Hamburger and Hamilton, 1951). In the domestic chicken, embryonic blood-derived PGCs can be propagated *in vitro* for several months, genetically modified and re-injected into early embryo surrogate hosts. Their ability to migrate to the gonadal *anlage* was unaffected by this treatment and the resulting hosts exhibited a high germline transmission rate (Van De Lavoir et al., 2006; Macdonald et al., 2010). For the zebra finch, it has been shown that primordial germ cells of the embryonic gonads (zfgPGCs) can be cultured for up to 30 days on a feeder cell layer in the presence of relative high concentrations of undefined fetal bovine serum, genetically modified with transposons and retain the ability to colonize embryonic host gonads (Jung et al., 2019). However, to the best of our knowledge the generation of transgenic songbirds using this approach has not yet been reported.

Here, we explored the use of blood-borne zfPGCs to generate transgenic zebra finches using a feeder-free culture medium that contained fetal bovine serum at a minimal concentration to avoid precocious differentiation and using lentiviral transduction for genetic modification. Cultured zfPGCs were efficiently transduced with lentiviral vectors expressing enhanced green fluorescent protein (eGFP). After injection underneath the early zebra finch blastodisc of a host embryo (EGK stage X), genetically modified zfPGCs migrated to and colonized the gonadal *anlage* following incubation. These surrogate host embryos were successfully hatched and raised by foster parents and, when mated, efficiently transmitted the transgene to their offspring (Fig. 1).

**Figure 1.**
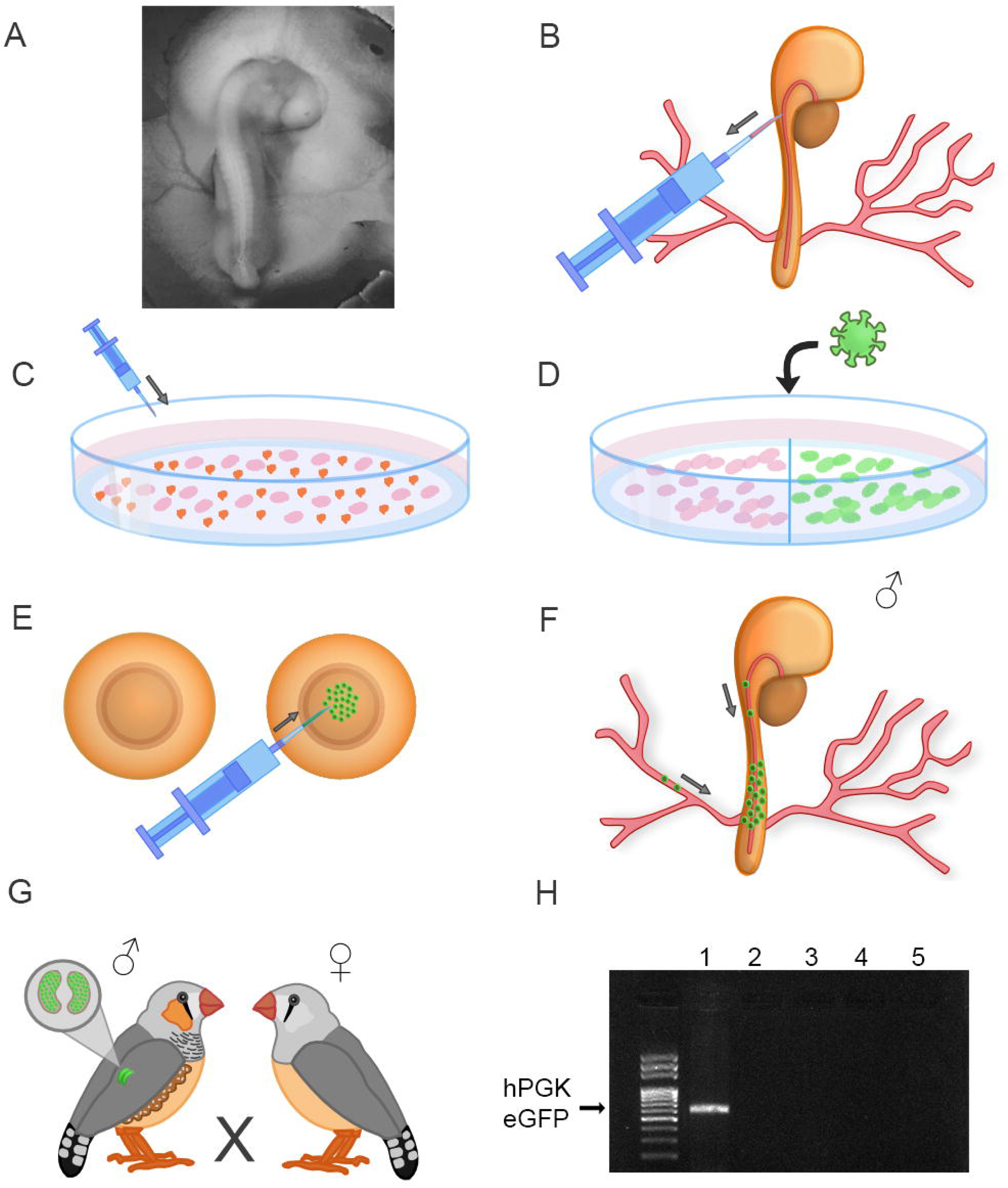
Method of transgenic zebra finch generation using cultured primordial germ cells. **(A)** Microphotograph of a zebra finch embryo at Murray stage 15. (B-G) Schematic representation of the procedure: **(B)** Blood extraction from the dorsal aorta of zebra finch embryos at Murray stage 13 to 15. **(C)** Selected expansion of zebra finch primordial germ cells (zfPGCs) in vitro. While erythrocytes started to die after 4 days in vitro (DIV), the proliferation of zfPGCs continued. **(D)** After 7 DIV, zfPGCs were transduced with lentiviral vectors for enhanced green fluorescent protein (eGFP) under control of the human phosphoglycerate kinase (hPGK) promoter. **(E)** zfPGCs that expressed eGFP at 10 DIV were harvested and injected into host embryos at the blastodisc stage. **(F)** During incubation of the injected eggs at 38°C for 60 hours eGFP-positive zfPGCs migrated through the developing circulatory system to the gonadal *anlage*. **(G)** Zebra finches with gonads that enclosed eGFP-expressing zfPGCs were raised to maturity and mated with wildtype birds. **(H)** The presence of the eGFP gene was determined by PCR of blood samples taken from the founders’ offspring. In blood samples that were taken from birds of the same clutch (1-5), transgenic offspring was identified by the presence of the hPGK-eGFP band (1).

## RESULTS

### Expansion of embryonic blood-derived zfPGCs in culture

Blood samples (1 - 3 μl) extracted from zebra finch embryos at Murray stages 13-15 (Murray et al., 2013) contained a mixed population of erythrocytes and about 140 PGCs, with large individual differences in the number of PGCs observed between samples (40 – 500). Like in chicken (Macdonald et al., 2010), blood-derived zfPGCs were round with a diameter of 16-20 μm, showed tiny membrane protrusions visible under phase contrast, a prominent nucleus and small cytoplasmic vesicles that enclosed polysaccharides which could be stained by the periodic acid-Schiff (PAS) reaction (Fig. 2 A-C).

**Figure 2.**
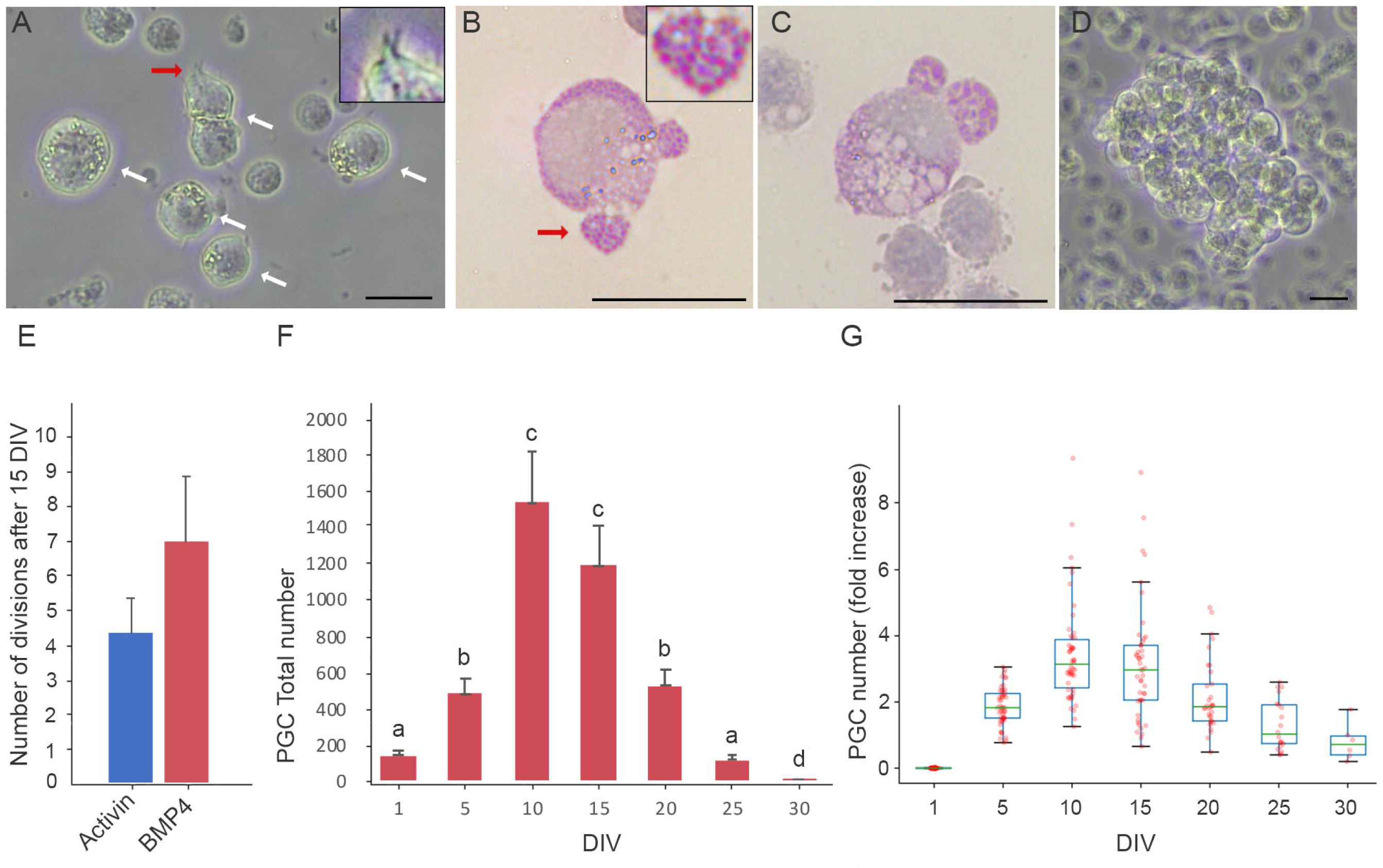
Characterization and expansion of cultured zfPGCs. **(A)** Phase contrast image of an early blood cell culture with zfPGCs (white arrows) that contained polysaccharide vesicles and showed typical cellular protrusions (red arrow). The insert in A shows cellular protrusions at higher magnification. Note that PGCs were larger in size than other blood cells. **(B, C)** Periodic acid-Schiff (PAS) stains of blood smear samples taken from a zebra finch **(B)** and chicken **(C)** embryo at embryonic stage 14, staining polysaccharide vesicles of PGCs in magenta. A cytoplasmic fragment of the zfPGC (red arrow) is shown in the insert at a higher magnification. **(D)** ZfPGCs growing in loose clumps at 7 DIV conserve their native morphology. **(E)** Blood samples from single embryos were split in two and cultured one half in the presence of Activin and the other with bone morphogenetic protein 4 (BMP4). Cell numbers were counted at 1 and 15 DIV and numbers of cell divisions at 15 DIV are shown (mean ± SEM). **(F)** Growth curve of zfPGCs cultured in medium containing BMP4. Different letters indicate statistically significant differences between cell numbers on different days (ANOVA; n=32, p=0.05). ZfPGCs proliferated significantly until 15 DIV. At 20 DIV a significant decrease of the cell number due to death was detectable. **(G)** Box plot showing fold increase in zfPGCs during a period of 30 days in culture as compared to the number of zfPGCs at 1 DIV. Scale bars in A - D represent 20 μm.

To culture zfPGCs, we used a feeder layer free culture and a defined culture medium (FAIcs) with low calcium concentration that we had developed for chicken PGCs (chPGCs) obtained from embryonic blood (Whyte et al., 2015), but replaced the chicken serum with a low concentration of fetal bovine serum. In this culture medium, we found that a low concentration of calcium (0.3 mM) supported the proliferation of zfPGCs in loose, non-adherent clumps (Fig.2 D, Fig.S1 A). Higher concentrations of calcium (medium with 100 % KnockOutTM DMEM containing 1.8 mM calcium) promoted stronger cell-cell interactions, leading to the formation of dense zfPGC clumps with indistinguishable cell boundaries. We also tested a culture medium previously developed for zfgPGCs (Jung et al., 2019) that contained higher serum concentrations. Using this medium condition, zfPGCs extracted from embryonic blood attached to the bottom of the culture plate (Fig. S1 B), showed a lower growth rate and exhibited signs of cell death.

To establish improved culture conditions for zfPGCs, we adapted the FAIcs by altering the ligand of the TGF-beta signaling pathway. Both growth factors Activin A and Bone morphogenetic protein (BMP) 4, have been shown to be sufficient for propagating chicken PGCs (chPGCs) *in vitro* (Whyte et al., 2015). However, we found a trend towards higher numbers of cell divisions in the presence of BMP4 for zfPGCs (average and SEM: 4.33 ± 0.88 for Activin and 6.97 ± 1.87 for BMP4; n=3; Fig. 2E), and replaced Activin with BMP4 for our germline transmission experiments.

When our optimized zfPGC culture medium was used (Table S1), proliferation of zfPGCs was detectable after 2 days *in vitro* (DIV), and loose cell clusters were observed forming after 5 DIV (Fig. 2D). During the first two weeks of culture, zfPGCs divided 2 to 9 times, reaching a maximum of 8000 cells. On average, we obtained 1331 zfPGCs at 10 DIV and 1045 zfPGCs at 15 DIV per blood sample (n = 32; Fig.2 F, G). Between 15 and 20 DIV, zfPGCs started to undergo cell death.

### Expression of cell type-specific markers by cultured zfPGCs

To test the effect of *in vitro* propagation on PGC-specific gene expression we assessed germ and stem cell marker expression by zfPGCs at 10 DIV by performing immunofluorescent stainings and RT-PCR analyses. Stage-specific embryonic antigen 1 (SSEA-1), a carbohydrate epitope associated with cell adhesion and migration, is normally expressed by avian PGCs during migration to the gonadal *anlage* in turkeys (D’Costa and Petitte, 1999), as well as in zfgPGCs (Jung et al., 2019) and also by chPGCs (Macdonald et al., 2010) *in vitro*. Furthermore, epithelial membrane antigen 1 (EMA-1), a cell surface glycoprotein, is expressed by chPGCs at early embryonic stages and *in vitro* (Urven et al., 1988; Raucci et al., 2015). Correspondingly, migrating zfPGCs that were freshly isolated from the embryonic blood stream at Murray stages 13-15 as well as zfPGCs at 10 DIV were found to be immunopositive for both SSEA-1 (Fig. 3A, D) and EMA-1 (Fig.3 E, H). In both antibody stainings, the cell surface of zfPGCs was strongly labeled (Fig. 3, D); in contrast, erythrocytes remained unstained (Fig. 3A, B, E, F).

**Figure 3.**
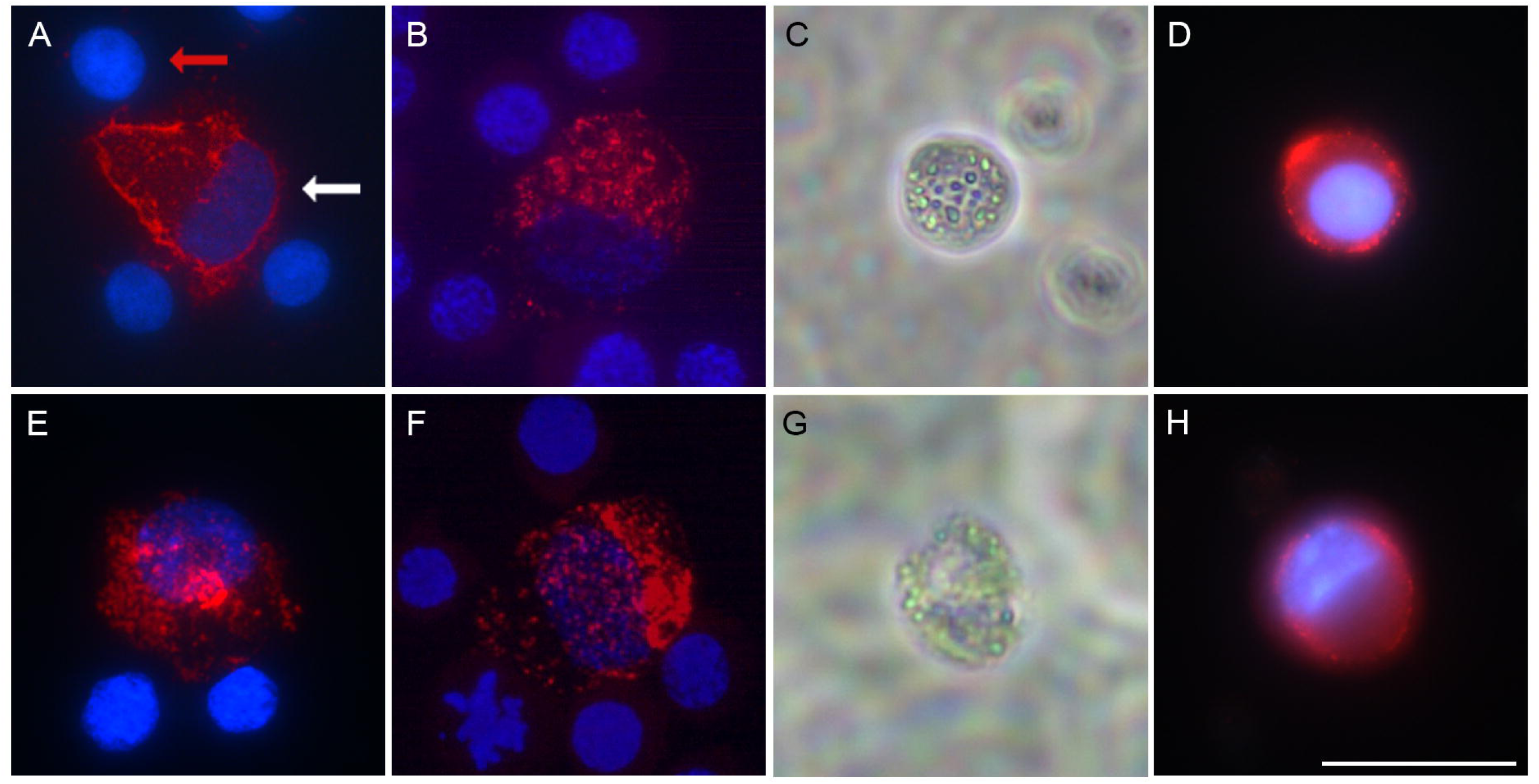
Immunostainings of zebra finch and chicken PGCs for cell type-specific markers. Microphotographs of PGCs were taken after immunofluorescent stainings for (**A - D**) stage-specific embryonic antigen 1 (SSEA1) and (**E - H**) epithelial membrane antigen 1 (EMA1). Stainings were performed with blood smears obtained from zebra finch (zf) (**A, E**) and chicken (ch) (**B, F**) embryos as well as with zfPGCs cultured for 10 DIV (**C, D, G, H**). Fluorescent images (immunostainings in red and nuclear stains with 4′,6-diamidin-2-phenylindol (DAPI) in blue) and phase contrast (**C, G**) images are shown. In (**A**) arrows point to a zfPGC (white arrow) and an erythrocyte (red arrow). Note that in blood smear samples PGCs were surrounded by immunonegative blood cells. Scale bar represents 20μm.

In both chPGCs and zfPGCs, the germline-specific genes DAZL (Deleted in Azoospermia Like) and DDX4 (DEAD-Box Helicase 4) as well as the pluripotency markers POU5F1 (POU Class 5 Homeobox 1) and NANOG (Nanog Homeobox) are expressed (Van De Lavoir et al., 2006; Macdonald et al., 2010; Jung et al., 2019). Our RT-PCR analyses revealed that cultured zfPGCs harvested after 10 DIV expressed these stem cell markers (Fig. 4). In control samples, zebra finch blastodiscs from freshly laid eggs, which contain PGCs, showed a similar expression pattern. In contrast, cultured fibroblasts from zebra finch embryos did not express these markers (Fig. 4).

**Figure 4.**
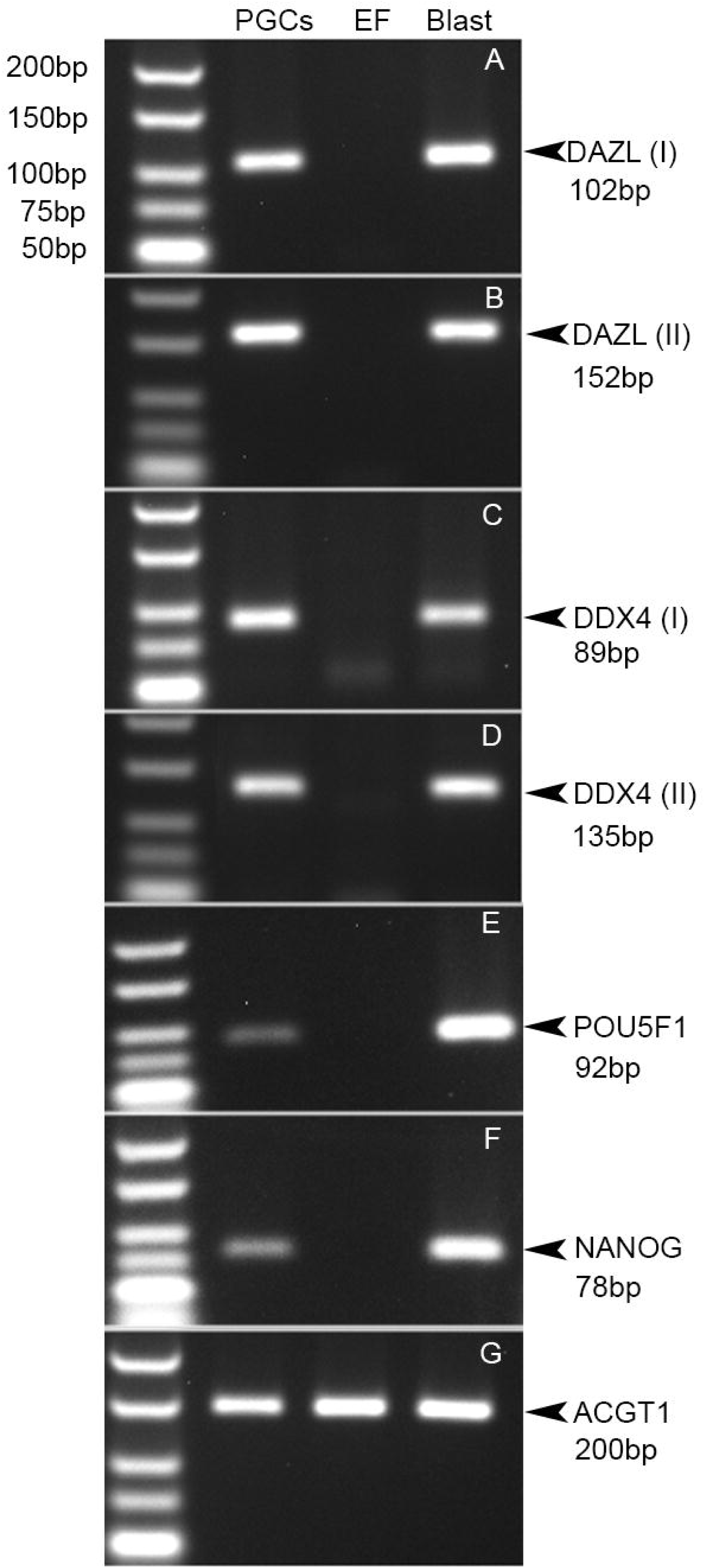
Conserved expression of germ and stem cell-specific marker genes in cultured zfPGCs. Photographs of agarose gels after electrophoretic separation and ethidium bromide staining of RT-PCR products. Cultured zfPGCs (PGCs; 10 DIV) continued to express germ cell markers such as DAZL **(A, B)** and DDX4 **(C, D)**, and, interestingly, also stem cell markers such as POU5F1 **(E)** and NANOG **(F)**. In contrast, cultured fibroblasts of zebra finch embryos did not express any of these marker genes. All cDNA samples from zebra finch blastodisc cells (Blast) that include early zfPGCs turned out to be PCR-positive for the same marker genes. Detection of y-actin (ACTG1) **(G)** expression was used to control all samples for equal cDNA quality. Sizes of molecular weight markers (left slots) and expected amplicons are indicated. For both DAZL and DDX4 two RT-PCRs were carried out using primer pairs that were reported by Mak et al. 2015 (I) and newly designed for this study to be intron-spanning (II) (see in material and methods).

### Highly efficient transduction of cultured zfPGCs with lentiviral vectors

Electroporation (BTX, Gemini System) and lipofection resulted in extensive cell mortality of cultured zfPGCs and low transfection efficiency (data not shown). To determine whether cultured zfPGCs could be genetically modified, we transduced cultured zfPGCs at 7 DIV with a lentiviral vector containing eGFP gene under control of the human phosphoglycerate kinase (hPGK) promoter (Fig. 5 A) when they were growing in cell clumps. Two days post-transduction, the majority of the cells exhibited strong eGFP expression (Fig. 5 B, C). In contrast to zfPGCs cultured from embryonic blood, zfPGCs cultured from embryonic gonads (zfgPGCs) (Jung et al., 2019) showed a relatively low lentiviral transduction efficiency (Fig. S1 G, H). We next compared the eGFP expression in cultured zfPGCs after transduction with lentiviral vectors that contained different constitutive promoters including the hUBC promoter, the human elongation factor 1 alpha (hEF1α) promoter, the cytomegalovirus (CMV) promoter and the CMV enhancer fused to the chicken beta-actin (CAG) promoter. For all these lentiviral constructs the level of the eGFP expression after transduction turned out to be similar (Fig. S2).

**Figure 5.**
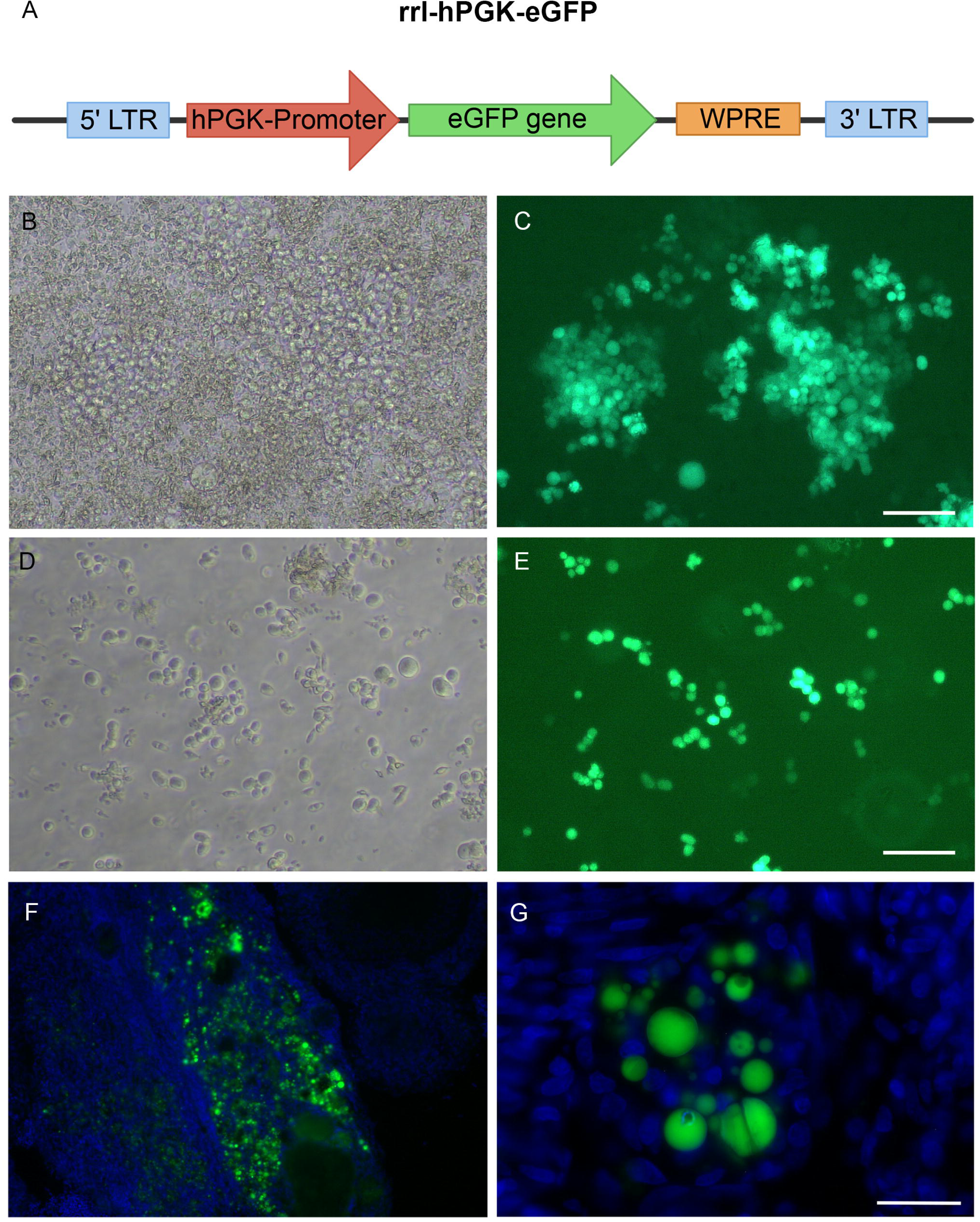
Cultured zfPGCs were efficiently transduced with a lentiviral vector for eGFP and colonized host embryo gonads. (**A**) Lentiviral vector rrl-hPGK-eGFP that was used to transduce zfPGCs *in vitro* and generate founder birds. **(B-E)** Microphotographs of cultured zfPGCs that were transduced with rrl-hPGK-eGFP for 48 hours. Phase contrast **(B, D)** and fluorescent images **(C, E)** are shown for zfPGCs expressing eGFP before **(B, C)** and after **(D, E)** dissociation of PGC clumps incubated with papain. Note that almost all zfPGCs were expressing the reporter gene. In **(F, G)** microphotographs of cryosections produced from an adult founder ovary that showed eGFP expression and blue nuclear counterstain with DAPI are presented at low (**F**) and high (**G**) magnification. Note that in the ovary eGFP-positive germ cells of different sizes display various stages of differentiation. Scale bar represents 100 μm.

Since viral titer is a crucial factor for a successful lentiviral vector transduction, we also assayed reporter gene expression in cultured zfPGCs when using increasingly diluted viral particle suspensions. Using the lentiviral vector for rrl-hPGK-eGFP, we found the highest eGFP expression at a final concentration of 2×10exp8 TU/ml and greater (Fig. S3). For gene editing studies in which Cas9, CRISPR gRNAs and a selection marker are used at the same time, it would be necessary to simoultaneously transduce zfPGCs with two different lentiviral vectors. To test this we treated the same cells with lentiviral vectors for both UBC-eGFP and CMV-Tomato at a titer of 2×10exp8 TU/ml each. Two days after transduction, most zfPGCs expressed both reporter genes at comparable strength (Fig. S4). Together, these findings demonstrated that lentiviral vectors provide an efficient and promising way to introduce transgenes into cultured zfPGCs.

### Upregulation of LDLR gene family members in cultured zfPGCs

The viral vectors used to transduce zfPGCs in this study were vesicular stomatitis virus glycoprotein G (VSV-G) pseudotyped lentivirus for which the low-density lipoprotein receptor family (LDLRF) constitutes the main cell surface receptors (Finkelshtein et al., 2013, Nikolic et al., 2018). Transcriptome analysis of zfPGCs that were freshly extracted from embryonic blood and cultured for 10 DIV revealed the expression of the LDLR gene family members LR11, LRP1B and LRP3 to be significantly up-regulated after 10 DIV. The expression of other gene family members such as LRP2, LRP6, LRP8 and LRP1 remained unchanged and was found to be down-regulated for VLDLR and LRP4 (Table 1). Absence of the prototypic LDLR gene in the zfPGC transcriptome was in line with the role of LDLR in steroidogenesis by somatic cells of chicken ovarian follicles (Hummel et al., 2003).

**Table 1.** Expression profile of Low density lipoprotein receptor family in zfPGCs cultured for 10 days

| Gene<br>Symbol | Gene Name | Zebra finch Ensembl<br>gene ID | Fold<br>Change | Adjusted<br>P. value |
| --- | --- | --- | --- | --- |
| LR11<br>(SORL1) | Low density lipoprotein<br>receptor relative with 11 ligand<br>binding repeats | ENSTGUG00000000462 | 2.816 | 0.05 |
| LRP1B | Low density lipoprotein<br>receptor related protein 1b | ENSTGUG000000011849 | 2.602 | 0,043 |
| LRP3 | Low density lipoprotein<br>receptor related protein 3 | ENSTGUG000000009422 | 1.8494 | 0.008 |
| LRP2<br>(Megalin) | Low density lipoprotein<br>receptor related protein 2 | ENSTGUG000000007663 | 1.172 | 0.175 |
| LRP6 | Low density lipoprotein<br>receptor related protein 6 | ENSTGUG000000012717 | 0.412 | 0.698 |
| LRP8<br>(APOER2) | Low density lipoprotein<br>receptor related protein 8 | ENSTGUG000000009223 | -0.008 | 0.994 |
| LRP1 | Low density lipoprotein<br>receptor related protein 1 | ENSTGUG000000015747 | -0.560 | 0.620 |
| VLDLR | Very low density lipoprotein<br>receptor | ENSTGUG000000005386 | -1.558 | 0,017 |
| LRP4<br>(MEGF7) | Low density lipoprotein<br>receptor related protein 4 | ENSTGUG000000010533 | -2.688 | 0,004 |

### Cultured zfPGCs colonized the gonadal *anlage* of host embryos after re-injection subgerminally

Cultured zfPGCs clumps expressing eGFP under control of the hPGK promoter were pooled from 5 to 10 cultures and dissociated after digestion with papain (Fig. 5 D, E). Around 500 cells were injected under the blastodisc of freshly laid zebra finch eggs. The eggs were sealed and incubated for 48 hours before being transferred into the nests of foster parents for further incubation. From 22 injected eggs 10 founder birds (45,4 %) were hatched and raised, six females and four males (Table 2). Using histological sections of founder gonads, we found that cultured and re-injected zfPGCs migrated to the gonadal anlage of the host embryo and differentiated into germinal cells (Fig. 5 F, G). Furthermore, integration of the hPGK-eGFP construct in genomic DNA extracted from founder gonads was verified by PCR in both ovaries and testes (Fig. S5 A). All founders that were examined contained hPGK-eGFP-positive zfPGCs in their gonads.

**Table 2.**
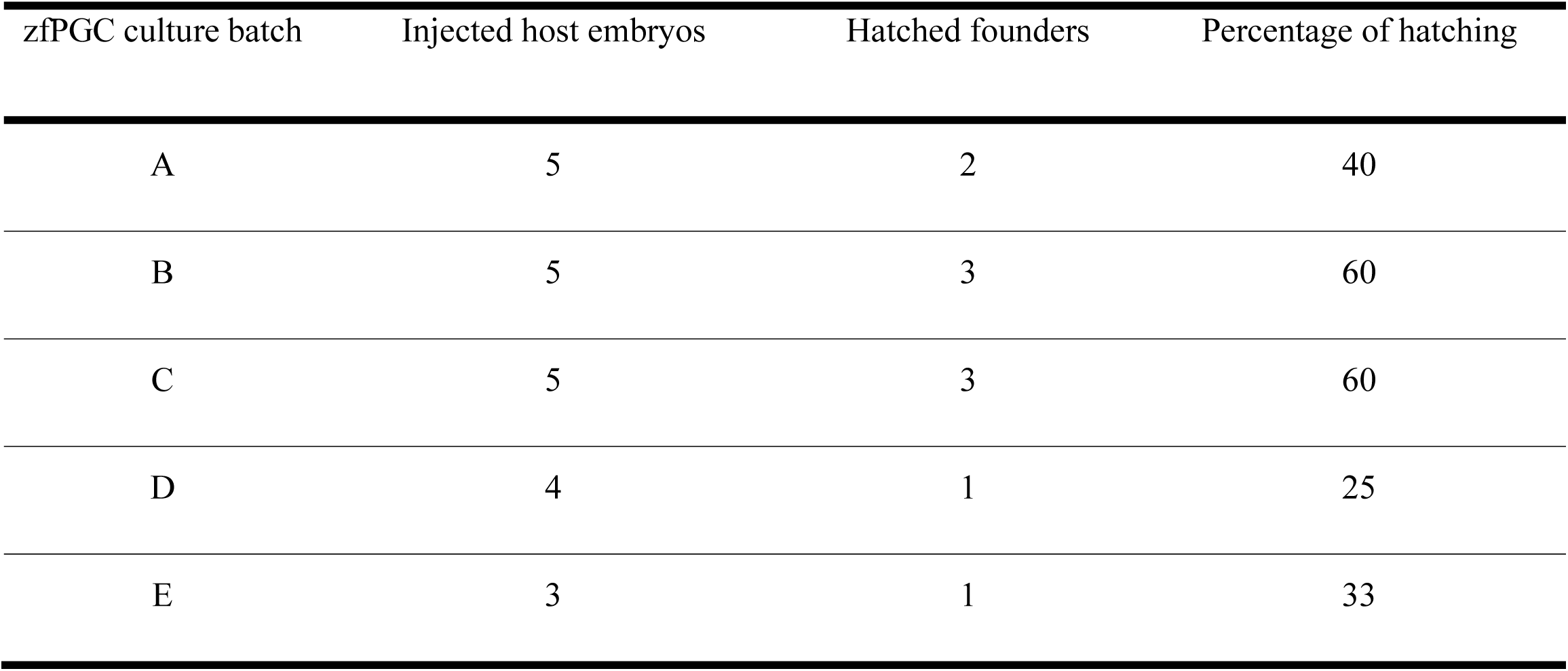
Hatching efficiencies of host embryos injected with cultured and transduced zfPGCs

### Songbird transgenesis

Founders were crossed with wild type birds, and genomic DNA from blood samples from the offspring (F1 birds) was analyzed by PCR for the presence of the hPGK-eGFP construct (Fig. S5 B). F1 birds that were hPGK-eGFP-positive by PCR were sacrificed to produce histological sections and confirm eGFP-protein expression by anti-GFP immunostainings (for control immunostainings of a wildtype zebra finch see Fig. S6). In transgenic F1 birds, we observed eGFP expression in the liver (Fig. 6 B) and GFP-immunopositive cells also in the brain, including forebrain song control nuclei like the HVC (Fig. 6 C, D), the robust nucleus of the arcopallium (RA; Fig. 6 E, F) and Area X. All F1 birds that were found to be hPGK-eGFP-positive by PCR turned out to be GFP-immunopositive as well. In summary, all 10 founder birds produced transgenic offspring showing a germline transmission rate between 4 and 22 % (Table 3). All transgenic birds generated where phenotypically normal and did not present any pathologies (Fig. 6 G).

**Table 3.** Frequency of germline transmission detected by PCR and anti-GFP immunostaining

| Founders |  | F1 birds |  |  |  |
| --- | --- | --- | --- | --- | --- |
| ID | Sex | Number | hPGK-eGFP-positive by<br>PCR | GFP-immunopositive | Germline<br>transmission (%) |
| 1 | Male | 18 | 4 | 4 | 22 |
| 2 | Male | 8 | 1 | 1 | 12 |
| 3 | Female | 9 | 1 | 1 | 11 |
| 4 | Female | 11 | 1 | 1 | 9 |
| 5 | Male | 23 | 2 | 2 | 8 |
| 6 | Male | 13 | 1 | 1 | 7 |
| 7 | Female | 17 | 1 | 1 | 5 |
| 8 | Female | 19 | 1 | 1 | 5 |
| 9 | Female | 20 | 1 | 1 | 5 |
| 10 | Female | 23 | 1 | 1 | 4 |

**Figure 6.**
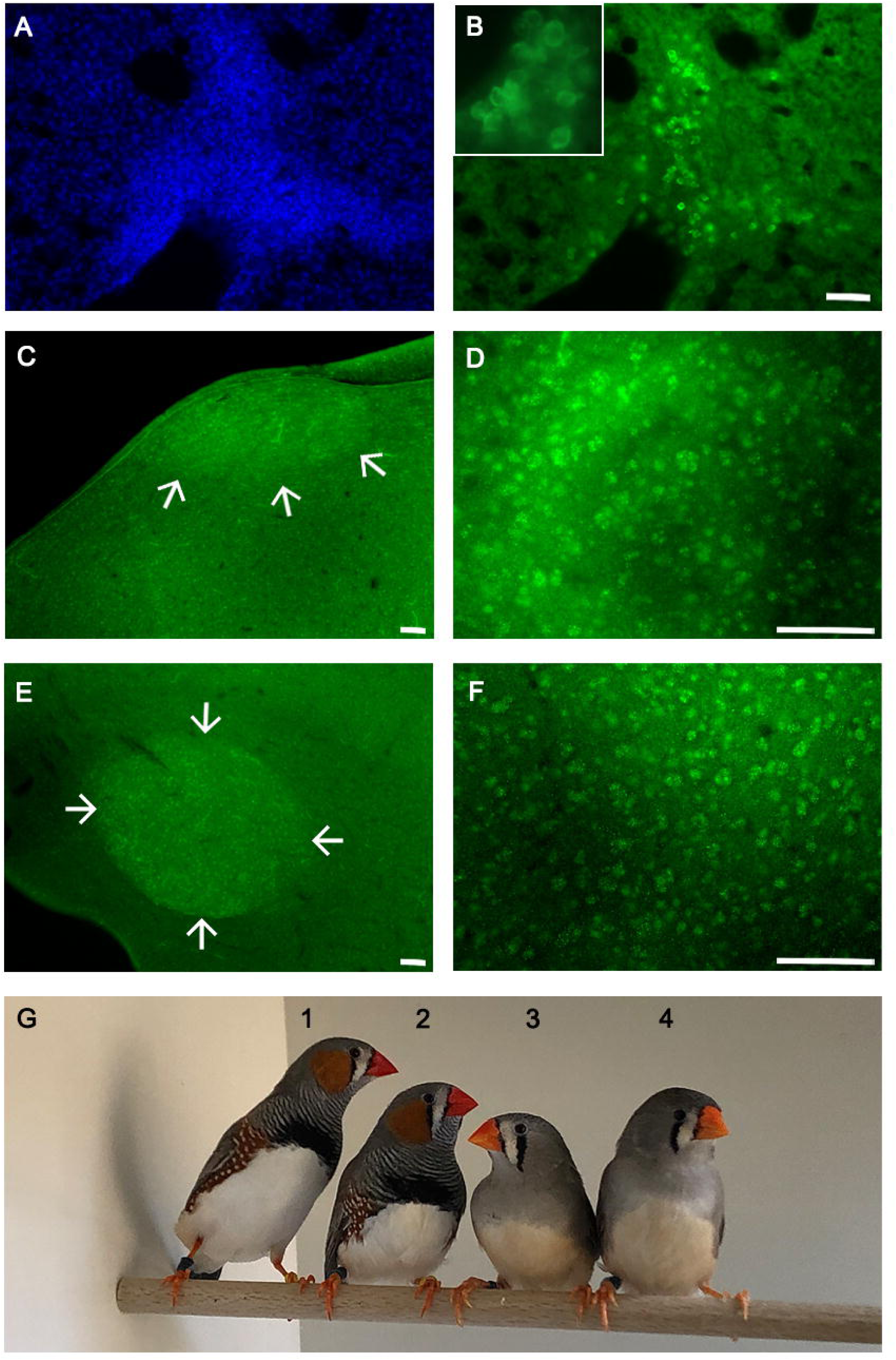
EGFP expression in liver and brain of transgenic birds. EGFP-expression in the liver (**A, B**) and GFP-immunopostive cells in the forebrain (**C – F**) of transgenic F1-birds. Liver was counterstained with DAPI (**A**) and insert in (**B**) shows eGFP-expressing liver cells at higher magnification. In (**C – F**) GFP-immunopositive cells are shown for male forebrain sections that included the song control nuclei HVC (**C, D**) and RA (**E, F**) at low (left panels) and high (right panels) magnification. At low magnifications location of the song control nucleus is indicated with arrows. (G) photograph of adult zebra finches 1, 3 being transgenic and 2, 4 being wild type. The scale bars represent 100 μm.

## DISCUSSION

Songbirds, especially the zebra finch are prominent animal models for studying neural circuit development, vocal communication and vocal learning among other topics (Bolhuis and Gahr, 2006; Konopka and Roberts, 2016). However, the wider use of the zebra finch in mechanistic research is hampered by the lack of transgenic technology, which is available for most sandard animal model organisms (Ménoret et al., 2017). In chicken, the long-term cultures of PGCs turned out to be extraordinarily useful for facilitating the production of transgenic chicken lines (Van De Lavoir et al., 2006; Macdonald et al., 2010). The ability to culture chPGCs almost indefinitely made it possible to modify the chicken genome *in vitro* and inject large numbers of genetically modified chPGCs into host embryos substantially increasing the germline transmission rate in these birds (Sid and Schusser, 2019; Schusser et al., 2013; Schusser et al., 2016; Dimitrov et al., 2016; Taylor et al., 2017; Oishi et al., 2016; Oischi et al., 2018). Here, we adapted such an approach for the zebra finch to allow for fast and efficient generation of transgenic songbirds.

We developed a new method for the propagation of zfPGCs from early embryonic blood cultures. When grown in cell clumps after 7 DIV, zfPGCs exhibited high transduction rates using lentiviral vectors such that almost every zfPGC expressed the reporter gene. Previously, the transduction and transmission efficiency of lentiviral vectors after injection at blastodermal stages was low (Agate et al., 2009). In particular, lentiviral vectors transduced zfPGCs much less efficiently *in ovo* compared to zfPGC clumps cultured *in vitro*. Factors like inactivation of lentiviral vectors in ovo and increased cellular uptake *in vitro* might be responsible for this difference. Thus, our approach constitutes a major improvement. We observed that lower transduction rates were detectable with singled zfPGCs obtained from cultured cell clumps after digestion with papain or prepared from embryonic gonads (Jung et al., 2019). Cellular interactions appear to promote the ability of lentiviral vectors to transduce the zfPGCs. Up-regulation of LDLR gene family members like LR11, LRP1B and LRP3 may have contributed to the increased uptake of the VSV-G pseudotyped lentiviral vectors by cultured zfPGCs. LR11 in particular, also participates in the cell adhesion process of hematopoietic stem cells by increasing the levels of urokinase type plasminogen activator (UPA) in the cell surface protoming plasmin formation (Nishii et al., 2013) and might aswell contribute to the formation of cellular aggregated in zfPGC. The highly efficient transduction of cultured zfPGCs with lentiviral vectors did not compromise the ability of the germ cells to differentiate in the host gonad and form functional gametes. All founder birds (100%) of both sexes that were generated produced transgenic offspring, which is astonishing considering the injected PGCs were derived from a mixture of male and female embryos. An additional factor that contributed to the high efficiency we achieved was the relatively high hatching success (45.4 %) of the injected host embryos. In comparison, injections of lentiviral vectors into blastoderm resulted in a hatching rate of 13.2% (Agate et al., 2009). We posit that the transfer of genetically modified zfPGCs into a host embryo by a single injection was less harmful to the embryo than the multiple vector injections required by the previous protocol used for the production of transgenic zebra finches (Agate et al., 2009). In conclusion, expansion of embryonic blood-derived zfPGCs *in vitro* followed by transduction with lentiviral vectors constitutes an extremely efficient strategy for producing transgenic zebra finches.

We selected lentiviral constructs containing the hPGK promoter to generate transgenic zebra finches because they generated consistently strong eGFP expression in cultured zfPGCs. In our transgenic hPGK-eGFP F1-birds, eGFP was not ubiquitously expressed in all zebra finch tissues but was detectable particularly in parts of the liver and after immunostaining in most brain areas including the song control system. The heterogeneous and selective expression pattern of eGFP in liver might be the result of a cell-type specific activity of the hPGK promoter in the zebra finch. For cell-type directed transgene expression, additional promoter constructs need to be tested to match the transgene expression pattern to the specific requirements that are defined by the various experimental questions to a transgenic songbird model.

The use of cultured zfPGCs that were transduced with lentiviral vectors appears to be an efficient transgenic approach for the overexpression of a transgene, and possibly, downregulation of an endogenous gene following random transgene insertion into the genome. The successful simultaneous transduction of cultured zfPGCs with two different lentiviral vectors suggested that in future CRISPR/Cas experiments the use of high-titer lentiviral preparations for Cas-proteins under the control of a PGK-promoter and gRNAs under control of an U6-promoter will be a promising approach. During a culturing period of two weeks we expect to be sufficiently long for the successful application of targeted gene editing techniques on zfPGCs, such as gene knock-outs by non-homologous end joining (NHEJ) repair of CRISPR/Cas-induced DNA breaks (Sid and Schusser, 2018). For more complex genome editing techniques that involve a homology directed repair (HDR), culture conditions that permit a more extended growth of zfPGCs in vitro and clonal selection of genetically modified zfPGCs will need to be developed. Nevertheless, for transgenic applications in neuroethological studies, the method of generating transgenic songbird models reported here represents a fast, efficient and straightforward procedure.

## MATERIAL AND METHODS

### Ethics statement

Animal handling was carried out in accordance with the European Communities Council Directive 2010/63 EU and legislation of the state of Upper Bavaria.

### Culture of embryonic blood-derived zfPGCs

Freshly laid zebra finch eggs were incubated for 60 hours at 37°C and relative humidity of 75% until Murray stage 13-15. Then a window was opened above the air chamber of the egg, and 1 - 3 μl of blood was extracted from the vasculature system using a pulled glass needle. Blood samples were cultured separately in a 96 well plate (Falcon) with 150 μl of the zfPGC culture medium in each well (Table S1). Cells were maintained in culture for up to 30 days and 80μl of the medium was replaced every other day. Gonadal PGCs (gPGCs) were obtained and cultured as described by Jung et al. (2019).

### Histological stainings

For periodic acid-Schiff (PAS) stain of blood smears, samples were extracted from zebra finch and chicken embryos at HH stage 14, dried on glass slides and fixed with 70% ethanol for 15 minutes. For staining, a PAS kit (Sigma Aldrich) was used according to the manufacturer’s instructions, and cells were counterstained with hematoxylin solution Gill No. 3 (Sigma Aldrich). For the immunofluorescence staining of freshly extracted zfPGCs, dried blood smears were fixed with 4% buffered formaldehyde for 10 minutes at room temperature. Slides were washed 3 times with washing buffer (0.2% Triton X100 in phosphate buffered saline (PBS) and blocked for 1 hour with a blocking solution of 10% pre-immune goat serum in washing buffer. After washing, samples were incubated for 12 hours at 4°C with a primary antibody against stage-specific embryonic antigen 1 (SSEA-1) (1µg/ml; Solter, D./Knowles, B.B.; DSHB Hybridoma Bank) or the primordial germ cell surface marker epithelial membrane antigen 1 (EMA-1) (5µg/ml; Eddy, M./Hahnel, A.; DSHB Hybridoma Bank) diluted in blocking solution. Thereafter, the slides were washed 3 times with washing buffer and incubated for 3 hours at room temperature with the secondary antibody (Alexa Fluor 594 goat-anti mouse (IgM); Life Technologies) diluted 1:300 in blocking solution. Following two washing steps with washing buffer, slides were incubated in 0.1μg/ml DAPI (4′,6-diamidino-2-phenylindole) diluted in water for nuclear staining, washed with PBS and mounted with 50% glycerol in PBS.

For cultured PGCs, the immunostaining was done in suspension with cells that we harvested after 10 days in vitro (DIV). Cells were centrifuged for 5 minutes at 2500 g, resuspended in 4% paraformaldehyde, incubated for 10 minutes at room temperature and washed twice with the washing buffer. Subsequently, washed cell pellets were resuspended and incubated in blocking solution for 1 hour at 20°C. After another washing step cells were resuspended in blocking solution containing the primary antibody and incubated for 12 hours at 4°C. Then, the cells were washed again and incubated with the secondary antibody for 3 hours at 20°C. Finally, cells were washed, stained with DAPI, washed again and resuspended in 50% glycerol in PBS. For immunostaining of living PGCs the fixation step was omitted. Images were obtained by performing epifluorescence microscopy.

### RT-PCR analyses

Total RNA was isolated from embryonic tissues using the RNeasy mini kit (Qiagen) and from cultured zfPGCs using the Power SYBR® Green Cells-to-CtT kit (Thermo Fisher), following the manufacturer’s instructions. For cDNA synthesis, 2 - 3 µg of total RNA were denaturated in 10 µl distilled water in the presence of 1µl random hexamers (50 ng/µl) and 1µl desoxyribonukleosidtriphosphates (dNTP mix; 10 mM each) for 5 minutes at 65°C. After chilling the solution on ice for 1 min, we added 2µl 10x reverse-transcriptase (RT) buffer, 4µl 25 mM MgCl2, 2µl 0.1 M 1,4-dithio-D-threitol (DTT), 1µl RNaseOUTTM (40U/µl; Thermo Fisher) and 1µl of the RT SuperScriptTM III (200U/µl; Thermo Fisher). Reverse transcription was carried out in a thermocycler for 10 min at 25°C, followed by 50 min at 50°C and, finally, 5 min at 85°C. Next, the reaction mix was placed on ice for 5 min and, after 1µl RNase H (2U/µl; Thermo Fisher) was added, incubated for 20 min at 37°C.

PCR reactions were carried out with the HOT FIREPol® DNA polymerase in buffer B1 (both Solis BioDyne). A 20 µl reaction mixture included 12.5µl H2O, 2µl 10x buffer B1, 1.6µl MgCl2 (25 mM), 1µl cDNA, 2µl primer pair (10 µM), 0.4 µl dNTP mix and 0.5µl HOT FIREPol®. The thermocycling conditions were as follows: denaturation at 95°C for 10 min, 35 amplification cycles (95°C for 30 sec, 60°C for 40 sec and 72°C for 30 sec) and a final extension at 72 °C for 5 min. The amplified products were resolved by gel electrophoresis in 2% Roti®agarose (Agarose High Resolution; Roth) in 0.5x UltraPureTM TRIS-Borat-EDTA (TBE) buffer (Thermo Fisher). Non-intron spanning primers for DAZL (DAZL(I)), DDX4 (DDX4(I)), NANOG, SOX3 and POU5F1 were the same as in Mak et al. (2015) (Mak et al., 2015). We designed the following intron-spanning primer pairs for zebra finch DAZL (DAZL(II): forward: 5’-GAAACCCAGCACTCAAACGC-3’; reverse: 5’-AAGACGCTCCGAATTTCAGC-3’) and DDX4 (DDX4(II): forward: 5’-CTGGAAGCCTACTCCAGTGC-3’; reverse: 5’-TCCCTCATCATTTGGGCCAC-3’). The primer pair designed for ACGT1 was forward: 5’-AACCGGACTGTTTCCAACAC-3’; reverse: 5’-CACCTTCACCGTTCCAGTTT-3’.

### Lentiviral transduction and injection of cultured zfPGCs

The lentiviral vector for eGFP under control of the human EF1α-promoter was purchased from SignaGen Laboratories (Gaithersburg MD, USA). All other lentiviral vectors were made in the department of Prof. Dr. Alexander Pfeifer (University of Bonn). For the production of lentiviral vectors, vector plasmids as well as the packaging plasmids pMDLg/pRRE, RSV-rev and pMD2.G were co-transfected into HEK293T cells seeded on poly-L-lysine-coated dishes. The supernatant was collected and centrifuged in an ultracentrifuge with SW32 Ti rotor at 61,700 g at 17 °C for 2 h. Virus suspensions were concentrated by ultracentrifugation over a 20% (w/v) sucrose cushion in a SW55 Ti rotor at 53,500 g at 17 °C for 2 h. Viral titer was quantified using reverse-transcriptase enzyme-linked immunosorbent assay.

To test for germline transmission, we used the vector for hPGK-eGFP. After 7 DIV, zfPGCs growing in clumps were pooled from 5 - 10 embryos without being separated by sex to reach a density of 1 – 3 x103 cells in 50μl of culture medium, and lentiviral vectors were added to a final concentration of 2 x108 TU/ml. The culture medium was replaced after 12 hours and the cells were cultured for additional 2 days. To dissociate zfPGC clumps, the cells were washed with PBS 48 hours after viral transduction and digested for 30 minutes at 37°C with a papain (Worthington, LS003119) solution (2mg/ml KnockOutTM DMEM without calcium) that had been sterilized using a 0.2μm syringe filter. Then the zfPGCs were washed with PBS, centrifuged for 5 minutes at 2,500 g and resuspended in culture medium to give a final concentration of 500 cells/µl for injection into a host embryo.

To inject transduced cultured zfPGCs freshly laid zebra finch eggs were incubated for 4 hours at 38 °C and then placed on a silicone surface with the blunt end facing upwards. With a light source that illuminated the egg from below, we were able to observe the blastodisc through the egg shell. Using a scalpel, we opened a 0.5-1 mm window in the egg shell above the blastodisc. Care was taken not to disturb the inner and outer egg membranes, as doing so would reduce the survival of the embryo. Using a pulled glass needle connected to a microinjector, we injected 300 - 500 transduced cultured zfPGCs in the subgerminal cavity of the host embryo. The window in the eggshell of the host embryo was sealed with chicken egg membrane and closed with zebra finch egg shell that had been glued to the host egg shell with zebra finch egg white. After 72 hours of incubation at 38 °C injected eggs that showed host embryo survival and ongoing development were taken to nests of foster parents for further egg incubation, hatching and development until sexual maturity.

### Generation and screening of transgenic zebra finches

Founder birds were raised by foster parents until sexual maturity and paired with wildtype birds. Fertilized eggs produced by the founders were incubated by foster parents that raised the hatchlings until maturity. In order to test for the presence of the eGFP gene in the offspring (F1-birds), blood samples were collected from each hatchling, and genomic DNA (gDNA) was extracted using the DNeasy Blood & Tissue Kit (Qiagen) following manufacturer’s instructions to perform PCR analyses. The primers used for the hPGK-eGFP-PCR produced an amplicon (602 bp) that included sequences of both the hPGK promotor and the eGFP gene: the forward primer was 5’-CACTAGTACCCTCGCAGACG-3’, the reverse primer was 5’-TCTTGTAGTTGCCGTCGTCC-3’. For each PCR 50 to 100 ng gDNA was used in a 20 µl reaction mixture that contained 12.5 µl H2O, 2 µl primer mix (10 µM), 0.4 µl dNTPs (10 µM), as well as 2 µl buffer B1 (10X), 1.6 µl MgCl2 (25 mM) and 0.5 µl FIREPol® DNA polymerase (all three from Solis BioDyne). Thermocycling conditions were the following: initial denaturation at 95°C for 5 min, 35 cycles of 95°C for 30 sec, 65°C for 30 sec and 72°C for 30 sec followed by 72°C for 5 min. The amplified products were resolved by electrophoresis in a 2 % Tris-acetate-EDTA (TBE) agarose gel. All steps had been performed in a lab free of eGFP usage or in an UV decontaminated PCR box.

To confirm eGFP expression by immunostaining, F1-birds that were PCR-positive for hPGK-eGFP were sacrificed and transcardially perfused with PBS followed by 4 % paraformaldehyde in PBS. Tissues were post-fixed in 4 % paraformaldehyde, cryoprotected in 10 % and 30 % sucrose in PBS, and sectioned at a freezing microtome. Free-floating tissue sections were washed three times for 10 minutes with PBS then incubated for 2 h in a PBS-blocking solution containing 0.5 % Saponin (Sigma Aldrich 84510) and 10 % pre-immune goat serum (Thermo Fisher). For anti-GFP immunostaining, sections were incubated for 36 h at 4°C with a chicken anti-GFP antibody (Aves GFP-1020) at a concentration of 1:1000 in PBS-blocking solution. After three subsequent washes with PBS, the samples were incubated for 3 h at room temperature with the goat anti-chicken antibody conjugated with Alexa Fluor® 488 (Abcam 150169) at a concentration of 1:500 in PBS-blocking buffer. Finally, the sections were washed three times with PBS and mounted on a glass slide with VECTASHIELD® (Vector laboratories). Images were obtained by performing epifluorescence microscopy. All birds that were PCR-positive for hPGK-eGFP and GFP-immunopositive were considered to be transgenic (Table 2). Gonads of founders were used freshly for DNA extraction followed by PCR for hPGK-eGFP and after fixation for anti-GFP immunostainings.

### Separation of cells by density gradient centrifugation

To separate zfPGCs from red blood cells two Ficoll 400 (Sigma F2637) solutions (16 and 6.3 %) were prepared in cell culture medium. Embryonic blood samples were pooled from five individuals and centrifugated for 5 min at 2,500 g. The cell pellet was resuspended in 100 μl of culture medium, mixed gently with 900 μl of the 16 % Ficoll solution, placed underneath 200 μl of the 6.3% Ficoll solution and centrifugated for 30 minutes at 3,400 g. ZfPGCs could be collected from the most superficial layer (200 μl) of the gradient. After 10 μl were removed for cell counting the rest of the cell suspension was diluted with 300 μl of culture medium and centrifugated for 5 min at 2,500 g. After the supernatant was carefully removed the cell pellet was resuspended in the amount of culture medium necessary for RNA extraction.

### RNA extraction and next generation sequencing

Four samples with 400 zfPGCs each were collected for each group (1 and 10 DIV) and RNA was extracted using the SMART-Seq v4 Ultra Low Input RNA Kit for Sequencing (TAKARA) following manufactures instructions. Illumina high output sequencing was done by the NGS group at the Max Planck Institute for molecular Genetics.

### Statistical analysis

For statistical comparison of zfPGC numbers obtained in cultures with BMP4 and Activin we performed a one-way ANOVA with p < 0.05. To analyze the growth of zfPGCs *in vitro* over time we performed a one-way ANOVA followed by a post-hoc test with the p-value being adjusted for multiple comparisons.

The Galaxy web platform (https://usegalaxy.eu/) was used for the analysis of the sequencing data (Afgan et al., 2018). A quality control of the FASTQ files was performed with FASTQC (http://www.bioinformatics.babraham.ac.uk/projects/fastqc/). Nextera adapters were removed with Trimmomatic (Bolger et al., 2014). The reads were mapped with STAR (Dobin et al., 2013) and counted with featureCounts (Liao et al., 2014). A quality control of the mapped sequences was done with MultiQC (Ewels et al., 2016). Differentially expressed genes between the two groups were identified with DESeq2 (Love et al., 2014).

## ACKNOWLEDGMENTS

We are grateful to Anja Lohrentz, Christina Reusch, Judith Kammerlander, Antje Bakker, Sunil Nandi and Lorna Taylor for their excellent technical support and Bernd Timmermann for performing the next generation sequencing of our samples. We want to thank David Witkowski, Frances Weigl and Frank Lehmann for their outstanding care of the birds in this study. Thanks go to Luisana Carballo for comments on the manuscript draft. IG also thanks the IMPRS research school for training and support. The study was founded by the Max Planck Society.

## AUTHOR CONTRIBUTIONS

I.G., F.D., M.H., M.M., M.G. conceived the study, M.G.,M.M. and A.P provided instruments, materials and reagents. I.G. performed the experiments. S.H. produced the lentiviral vectors. CFV contributed to the analysis of the sequencing data I.G. wrote the manuscript and all coauthors contributed to manuscript revision. I.G. is a member of the International Max Planck Research School (IMPRS) for Organismal Biology and the work was funded by the Max Planck Society.

