## Supplemental for "Highly efficient genome modification of cultured primordial germ cells with lentiviral vectors to generate transgenic songbirds"

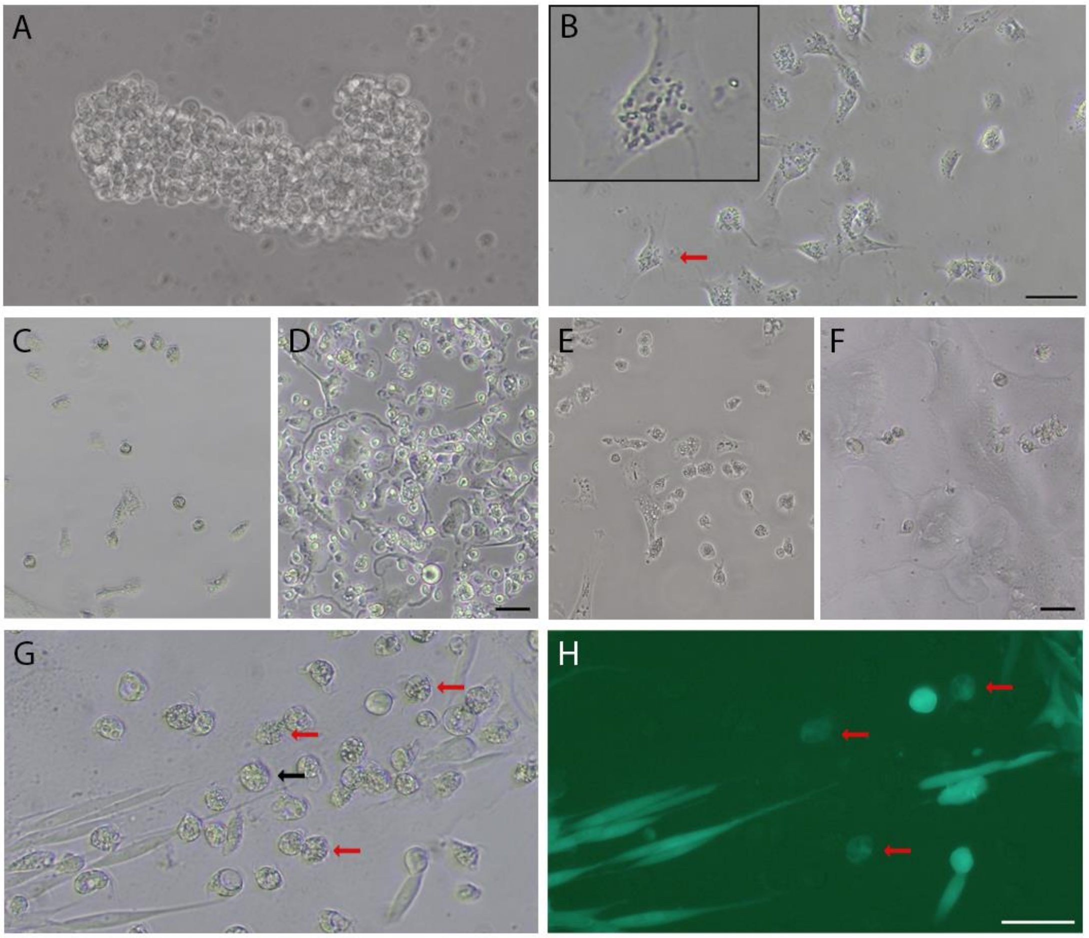


**Figure S1. Comparisons of culture conditions for zfPGCs extracted from embryonic blood and gonads**

Phase contrast images are shown for embryonic blood-derived zfPGCs cultured for 10 DIV in **(A)** the adapted FAIcs culture medium (Tab. S1) and in **(B)** gonadal PGC culture medium (Jung et al., 2019). The insert in (**B**) shows a zfPGC (red arrow) at higher magnification. Note that embryonic blood-derived zfPGCs cultured in adapted FAIcs conserved a rounded shape when growing in clumps (**A**) but attached to the bottom of the cell culture plate, changed their morphology and died shortly after when cultured in the gonadal PGC culture medium (**B**). In **(C - F)** phase contrast images are presented for zfPGCs that were extracted from embryonic gonads (zfgPGCs) and cultured in gonadal PGC culture medium (**C, D**) and adapted FAIcs (**E, F**), respectively. Images were taken after 1 (**C, E**) and 10 (**D, F**) DIV, respectively. Note that after 10 DIV in gonadal PGC culture medium zfgPGCs have proliferated and a feeder layer was formed with stromal cells (**D**) whereas in adapted FAIcs the zfgPGCs died (**F**). However, zfgPGCs were transduced with a lentiviral vector at relative low efficiency even when cultured in gonadal PGC culture medium. Phase contrast and **(G)** fluorescent **(H)** images of zfgPCRs are shown after transduction with a lentiviral vector for eGFP under control of the human UBC (hUBC) promoter. In (**G, H**) red arrows point to successfully transduced zfGPGCs and in (**G**) the black arrow hint at one of the zfGPGCs that did not express eGFP.

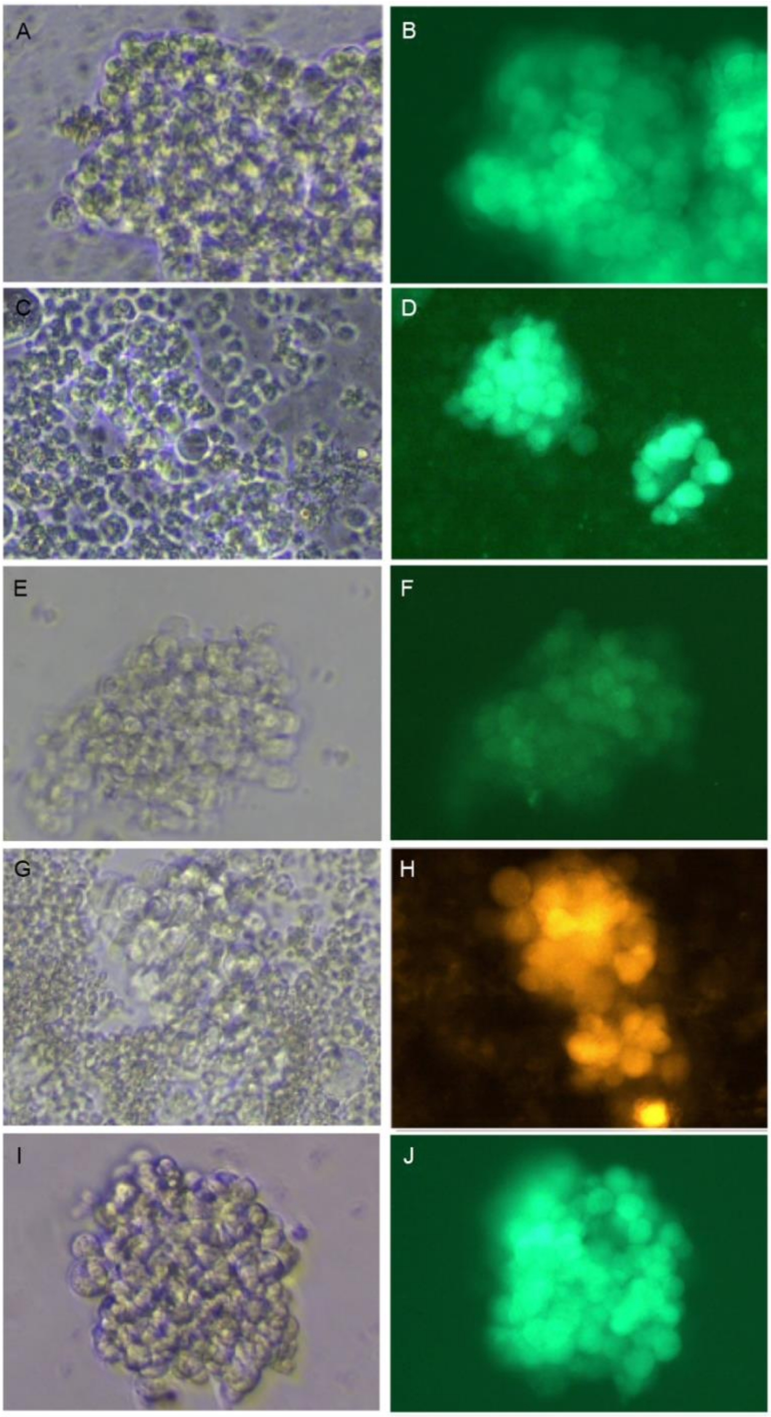


**Figure S2. Lentiviral vectors containing different promoters were capable of driving the reporter gene expression in cultured zfPGCs**

Phase contrast (left panels) and fluorescent (right panels) images of cultured zfPGC clumps that were transduced with lentiviral vectors for eGFP (**A - D, I, J**) and tomato (**E, F**). Lentiviral constructs contained different promoters: **(A, B)** human phosphoglycerate kinase (hPGK) promoter, **(C, D)** human ubiquitin C (hUBC) promoter, **(E, F)** human elongation factor 1 alpha (hEF1α) promoter, **(G, H)** cytomegalus virus (CMV) promoter and **(I, J)** CMV enhancer fused to the chicken beta-actin (CAG) promoter. The titer we used was 2x10exp8 TU/ml. Scale bar represents 20 μm.

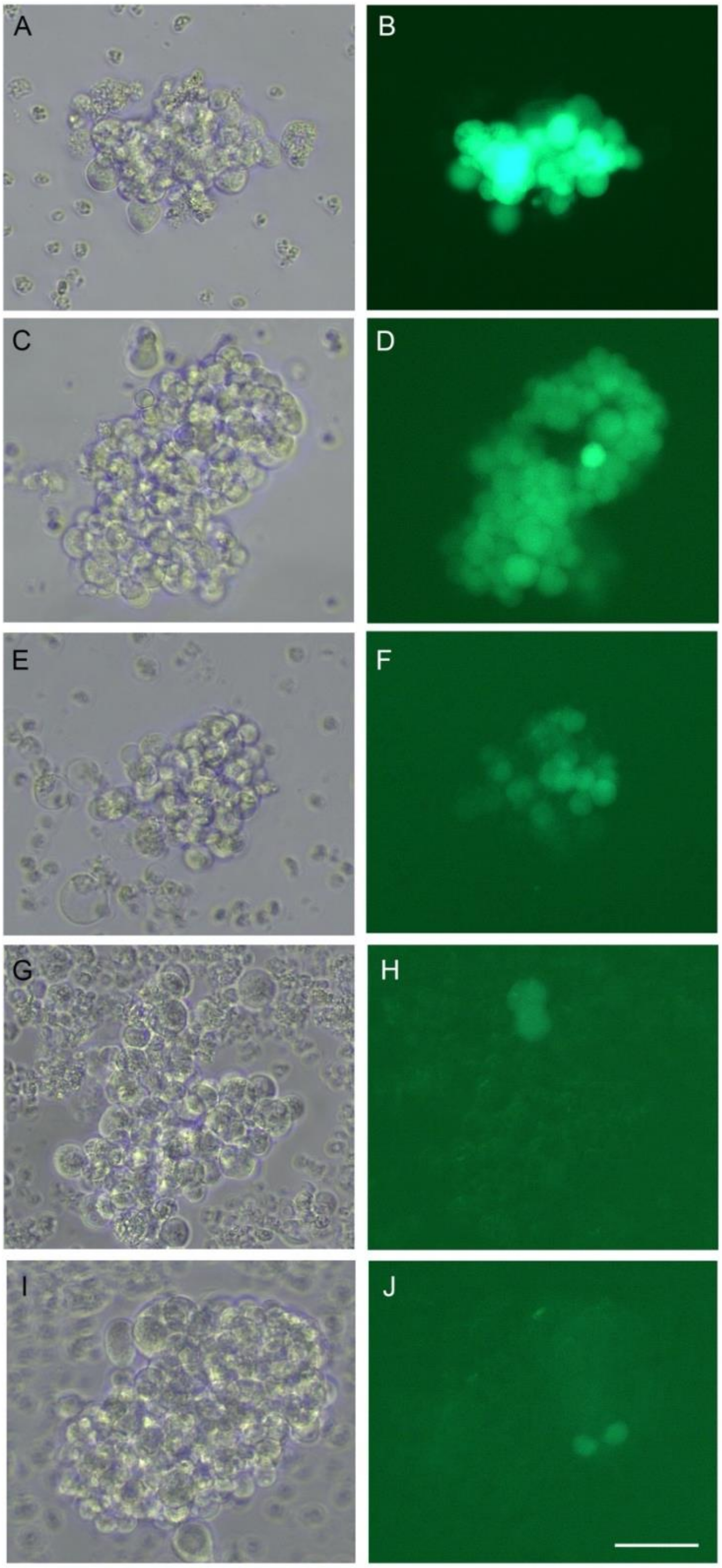


**Figure S3. Determination of effective lentiviral titers for the transduction of cultured zfPGCs**

Phase contrast (left panels) and fluorescent (right panels) images of cultured zfPGC clumps that were transduced with the lentiviral vector rrl-hPGK-eGFP using different viral titers (final concentrations): 4x10exp8/ml TU/ml **(A, B)**, 2x10exp8 TU/ml **(C, D)**, 4x10exp7 TU/ml **(E, F)**, 2.7x10exp7 TU/ml **(G, H)** and 8x10exp6 TU/ml **(I, J)**. Note strong expression of eGFP at a concentration of 2x10exp8 TU/ml and above. Scale bar represents 50 μm.

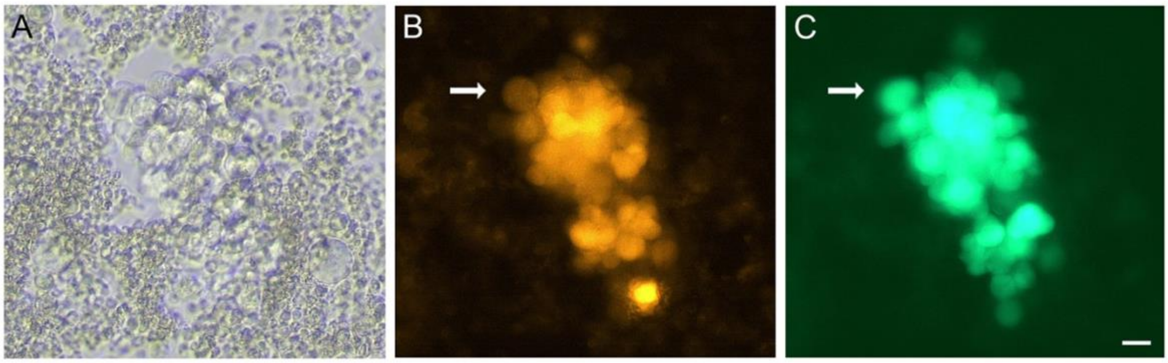


**Figure S4. Transduction of cultured zfPGCs with two different lentiviral vectors**

Microphotographs of cultured zfPGC clumps that were simultaneously transduced with lentiviral vectors for hUBC-eGFP and CMV-tomato. Phase contrast image **(A)** as well as fluorescent images of eGFP **(B)** and tomato **(C)**, are shown. White arrows point to the same zfPGC being fluorescent with both filters. Note that most zfPGCs expressed both eGFP and tomato. Scale bar represents 20 μm.

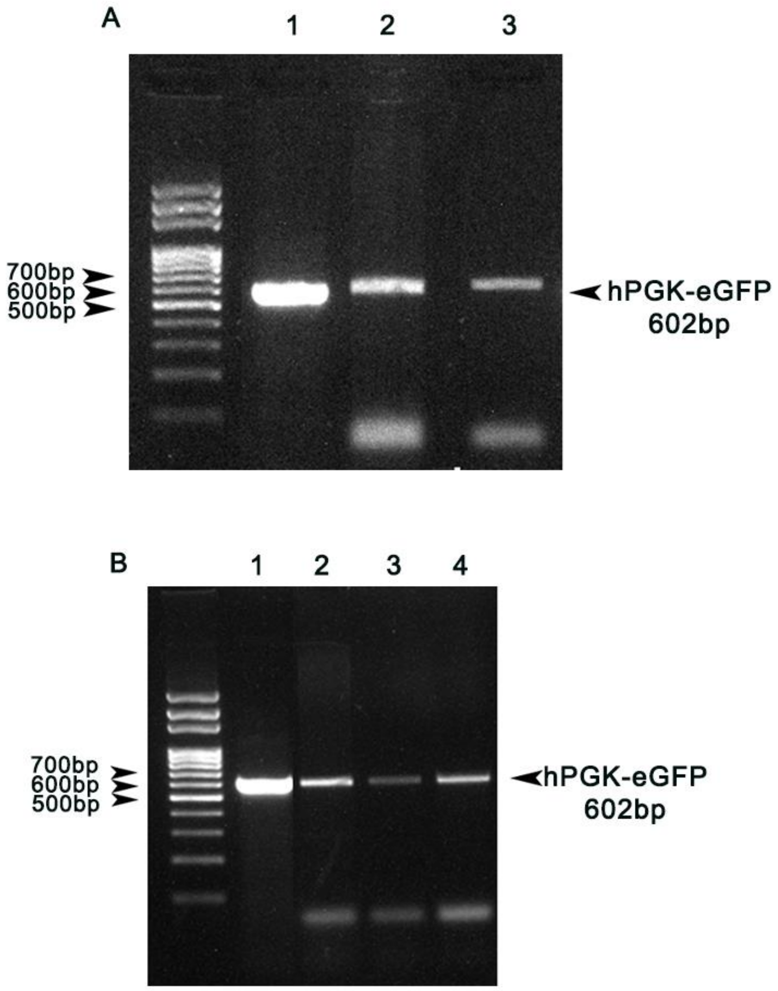


**Figure S5. Detection of the lentiviral construct in genomic DNA samples by PCR**

Genomic DNA (gDNA) was isolated from **(A)** founder gonads and **(B)** blood samples of transgenic F1 birds to detect a hPGK-eGFP sequence (602 bp) of the lentiviral construct by PCR. After electrophoretic separation images of agarose gels are shown that contained a molecular weight marker (left lanes) and PCR products that were obtained using the lentiviral transgene plasmid (**A**, 1; **B**, 1), gDNA isolated from a founder ovary (**A**, 2) and testis (**A**, 3), and blood gDNA of three F1-birds, two males (**B**, 2 and 3) and one female (**B**, 4).

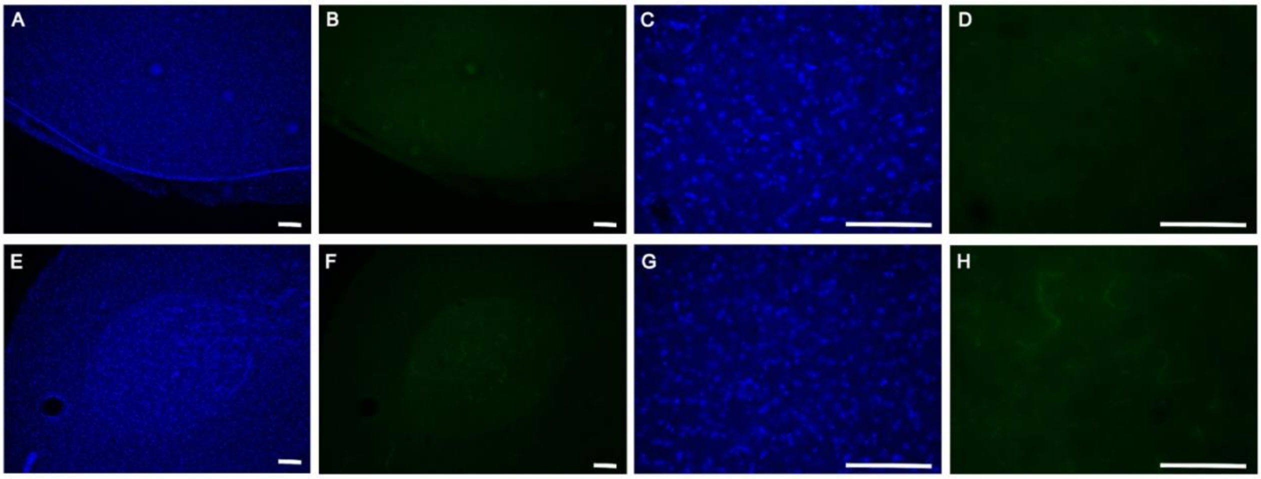


**Figure S6. Anti-GFP-immunostainings of control brain sections.**

Nuclei stains with DAPI **(A, C, E, G)** and anti-GFP immunostainings **(B, D, F, H)** were performed with brain sections obtained from a wildtype male zebra finch. Microphotographs of fluorescent images are shown for song control nuclei HVC (**A - D**) and RA (**E - D**). In (**C, D, G, H**) higher magnifications are shown for images presented in (**A, B, E, F**), respectively.

### Supplementary Table 1. FAIcs culture medium adapted to culture zfPGCs

| Culture medium components | | Concentration | Brand | Catalog number |
|---|---|---|---|---|
| Basal medium | KnockOut™ DMEM without Calcium | 70% | Thermo Fischer Scientific (TFS) | Custom made |
| | KnockOut™ DMEM with Calcium | 30% | TFS | 10829018 |
| Supplements | B27 Supplement | 1,5% | TFS | 17504044 |
| | Ovalbumin | 1% | Sigma Aldrich | A-5503 |
| | GlutaMax | 1% | TFS | 35050061 |
| | Non-Essential Amino Acids | 1% | TFS | 11140-050 |
| | Nucleosides | 1% | Sigma Aldrich | ES-008-D |
| | Fetal Bovine Serum | 0.4% | Sigma Aldrich | 12103-C |
| | Sodium Heparin | 0.2% | Sigma Aldrich | H3149 |
| | β-Mercaptoethanol | 0.25 mM | TFS | 31350-010 |
| | Pyruvate | 0.2 mM | TFS | 11360-039 |
| | Cholesterol | 4000 ng/ml | Sigma Aldrich | C3045-5G |
| | Human BMP4 | 25 ng/ml | TFS | PHC9534 |
| | Human FGF2 | 5ng/ml | R&D systems | 234-FSE-025 |
| | Human IGF | 25ng/ml | R&D systems | 291-G1-200 |
